# Distinct Neurodevelopmental Patterns and Intermediate Integration-Based Predictive Modeling in Autism Spectral Disorder

**DOI:** 10.1101/2023.06.26.546426

**Authors:** Yanlin Wang, Shiqiang Ma, Ruimin Ma, Linxia Xiao, Shi Tang, Yanjie Wei, Yi Pan

## Abstract

Autism Spectrum Disorder (ASD) is known to exhibit a more rapid expansion in brain structure during the early years of life compared to typically developing children (TD). This is of utmost importance in understanding atypical brain development and forecasting the onset of ASD. However, the precise age-related cortical changes (trajectories) that could pinpoint atypical brain development in ASD remain largely unknown. In this study, we characterize the distinct developmental patterns of cortical morphology in individuals with ASD, investigate the important neural biomarkers for ASD diagnostics, and propose a deep-learning workflow that combined graph convolutional networks with low-rank multi-model tensor fusion (LMFGCN) for ASD prediction. Our findings reveal that the constituents of gray matter volume (GV), cortical thickness (CT), and surface area (SA) exhibit separable developmental trajectories in ASD. Furthermore, we identify regional differences in CT and SA that underscore the separable brain developmental trajectories, both of which contribute to changes in GV. Our study also demonstrates that LMFGCN, an end-to-end deep-learning model with intermediate integrative approach, outperforms early and late integration methods and other state-of-the-art models in ASD classification. Overall, our results highlight the importance of distinguishing between cortical SA and CT for understanding ASD pathobiology, particularly during the early brain overgrowth period, and demonstrate the potential utility of LMFGCN in ASD classification.

## INTRODUCTION

Autism spectrum disorder (ASD) is an early-onset neurodevelopmental disorder characterized by early impairments in social communication and interaction, as well as restricted, repetitive patterns of behaviors, interests, or activities [1, 2]. These core symptoms typically manifest from around age 2 years and are accompanied by developmental differences in brain anatomy, functioning, and connectivity that affect behaviour across the lifespan [3-5]. While the etiology of autism remains largely unknown, it has been suggested that genetic, developmental, and environmental factors might act either alone or in combination as possible causal or predisposing effects toward developing autism [6]. The neuropathological mechanisms underlying ASD seem to revolve around disrupted molecular and cellular processes that lead to cortical abnormalities in infants with typical developmental controls (TD) [7, 8]. Altered maturational rates within different brain regions during early development may result in structurally and functionally distinct architecture throughout life [9, 10]. As such, a substantial normative dataset characterising the developmental trajectory of different morphometric features cross the lifespan, would be imperative in achieving a comprehensive understanding of ASD brain development. Moreover, novel data analytical and integrated approaches are crucial in facilitating the identification of anatomical abnormalities in individuals with ASD.

Several magnetic resonance imaging (MRI) studies have focused on the role of neurodevelopment trajectory of brain maturation compared to TD during early childhood that might been implicated in ASD [11-13]. Note that the atypical pattern of brain development in individuals with ASD is not linear across different stages of neurodevelopment [14-16]. For example, bran volume overgrowth (gray and matter volume) in ASD commonly occurs in the first 2 years of life [17-19] and followed by a plateaued growth [13, 20] and possibly a decrease in brain volume over adolescence [5, 21], which temporally correlates with postmortem findings of increased synaptic pruning during adolescence and early adulthood [22-24]. Hazlett and colleges [17, 19] prospectively reported a generalized cerebral cortical enlargement, driven by an increase in brain surface area but not cortical thickness, in very young children with ASD, which is important for understanding of cellular and genetic mechanisms. However, the direction and regional sequence of microstructural growth remain controversial. Other studies [9, 21, 25, 26] have reported that the neurodevelopmental trajectory of the brain morphological changes in ASD varies across different brain regions, indicating a disruption of the typical early brain development sequence (i.e., from back to front). Hence, elucidating the neuropathological mechanisms that underlie ASD necessitates a thorough comprehension of these brain temporal and regional alterations [13, 19, 27].

Furthermore, to fully utilize multiple types of features for predicting individuals with ASD, advanced deep learning models and data integration methods has gained popularity in medical imaging applications [28]. For instance, Parisot et al. [29] utilized graph convolutional networks (GCNs) to merge MRI data and pairwise interactions using phenotypic data by executing node classifications on a population graph for ASD classification. Alternatively, Arya et al. [30] employed relational information from sMRI data in comparison to phenotypic data alongside multiple MRI data for ASD classification utilizing GCNs. Additionally, Wang et al. [31] presented a multi-atlas graph convolutional network method that was based on varied brain atlases for identifying ASD patients and then combined different feature representations using a stacking ensemble learning method. However, these existing GCNs methods primarily focus on semi-supervised methods by propagating the labels from labeled data to unlabeled ones, which cannot be directly applied to predicting new samples [32-35]. Moreover, while current GCNs-based methods often implement early and late integration strategies to combine diverse types of data [36-38]. These models cannot leverage distinct (intra-modality) and complementary (inter-modality) information between different modalities. Unlike early and late integration, we proposed a novel intermediate integration method that can effectively extract more informative representations from each modality while also exploiting the interplay of information between intermediate representations from data transformation techniques [28, 39]. Moreover, we compared our model with early and late integration strategies as well as traditional machine learning methods to evaluate the learning performance for ASD classification.

Overall, combining individual subject data from KQJH, 16 cohorts were collected previously as part of the ABIDE consortium [40], and 17 as part of ABIDE-II [41], with a focus on three domains: brain developmental trajectory, neural biomarkers, and advanced disease classification model. Our initial approach involved adopting the normative modelling methods to analyses the different normative trajectories of brain morphometric properties for ASD and TD over the lifespan. Subsequently, we investigated important neural biomarkers from multiple types of cortical features related to the disease. Additionally, we utilized MTFGCN, an end-to-end deep-learning model with intermediate integrative approach, for ASD classification. Altogether, our study endeavors to achieve three primary objectives: firstly, to identify atypical brain development trajectory in ASD comparing with TD across the lifespan, which may point towards specific neurobiological mechanisms for ASD. Secondly, to get insights into the neural substrates underlying the ASD phenotype, which provide crucial clues to help in the development of new treatments. Finally, to build an advanced deep-learning workflow for ASD classification, which has the potential to enhance ASD prevention and treatment outcomes.

## METHODS

### Participants

All data are acquired from ABIDE-I [40], ABIDE-II [41] and KQJH. ABIDE-I consists of 1112 participants in total, 539 ASD participants and 573 TD participants. ABIDE-I data was collected across 17 scanning sites. Ages (in years) ranged between 7-64 in the ASD group and between 6-56 in the TD group. ABIDE-II consists of 1114 participants, 521 ASD participants and 593 TD participants. ABIDE-II data was collected across 19 scanning sites. Ages (in years) ranged between 5-62 in the ASD group and between 5-64 in the TD group. KQJH dataset is originally collected to investigate the association between genetic, neuroimaging and clinical data for children with ASD. All data was collected in the same site. The MRI dataset currently consists of 323 participants in total, 237 ASD participants, 86 TD participants. Ages (in years) ranged between 0-7 in the ASD group and between 0-4 in the TD group. All datasets utilized in this study were obtained from research that was authorized by local Internal Review Boards and conducted in compliance with pertinent ethical guidelines and regulations. Additionally, informed consent was obtained from all participants, or in the case of minors (under 18), consent was obtained from their parent or legal guardian.

### MRI acquisition and preprocessing

Due to the rapid brain development in childhood, MRI preprocessing pipeline is not well developed for this specific stage. To deal with this challenge, we used different preprocessing steps for different age stages. For 0-2-year-old children, we used the month-wised MRI preprocessing by Infantfreesurfer tools [42]. For 3-6-year-old children, we first applied the age-specific brain template and created the prior masks using ANTs [43]. Then, we conducted age-wised preprocessing using Freesurfer [44, 45]. For participants whose age over 6 years old, we used a standard MRI preprocessing pipeline in Freesurfer [44]. All details of MRI acquisition and preprocessing process are summarized in the Supplementary Material. The FreeSurfer algorithm automatically parcellates the cortex and assigns a neuroanatomical label to each location on a cortical surface model based on probabilistic information. The parcellation scheme of the Desikan–Killiany atlas [46, 47] was used to divide the cortex into 34 regions per hemisphere. For each participant, left and right cortical thickness (CT), surface area (SA), and gray matter volume (GV) measures were calculated.

### Quality assurance (QA) and exclusion criteria

The quality control was assessed by inspecting the image and the cortical reconstructed results across individual brains by two independent reviewers (YL.W. and S.T.). To enhance the accuracy of QA and to investigate the reasons of image exclusion, we also use quality control software for MRI data (MRIQC, [48]) to compute image quality metrics for each dataset (see Table. S8/9/10). In case of disagreement, the participant was discarded if any of the independent reviewers indicated the issue of image. Detail information for QA and exclusion criteria, please supplementary materials.

### Modeling developmental trajectories of brain morphology

The brain in infants with autism undergoes an abnormal morphological and volumetric changes compared to TD, contributing to clinical abnormalities in behavior and cognitive function. The approach of estimating mean and standard deviation from the residuals won’t work for brain development over the lifespan, such as heteroskedastic and non-gaussian models [49-52]. Generalized additive models for location, scale and shape (GAMLSS) can be used to model multi-parameter distributions, where each parameter of the distribution, i.e., location, scale, and shape, can be modeled as a function of several input variables, thereby accommodating a wide range of distributions [53-55]. So, for a four-parameter distribution:

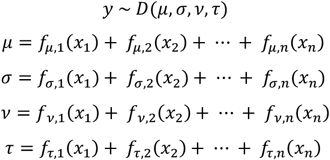

Where y is the response variable; *x*_*n*_ are subsets of the exploratory variable; *D*() is a probability distribution with parameters location (*μ*), scale (*σ*), skewness (*v*), and kurtosis (*τ*); *f*(*x*_*n*_) is a smoothing function of*x*_*n*_ for each parameter.

To capture the developmental trajectory of ASD and TD, GAMLSS models were fitted to structural MRI data for the mean CT (MCT), total SA (TSA), and total GV (TGV) separately. First, all data was split into 70% train dataset and 30% test dataset. Given the effect of age over the lifespan seems to be slightly non-linear and non-gaussian distribution, Sinh-Arcsinh (SHASH) [56] was adopted to estimate a model for four parameter distribution. We next compare several models with different additive items for ASD and TD group: We can only choose model age smooth effect for μ and σ (M1a), and add sex and site intercepts for μ and σ as a fixed effect (M1b), and add age smooth effect for *v* and *τ* (M1c), and add sex and site intercepts for *v* and *τ* (M1d), and adjust sex and site for μ and σ as random effects (M1e).

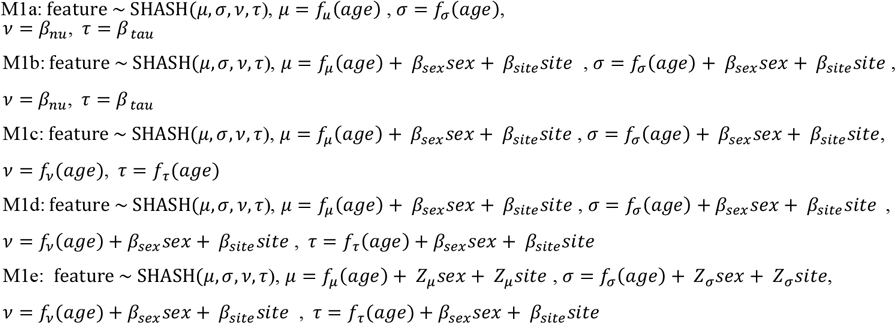

The best model was selected by Akaike Information Criterion (AIC) and Bayesian information criterion (BIC) for all solutions. Then, we compared each models in the training set and then validate the models in the testing set. Finally, we used the *mlr* package to evaluate the performance of 5 models. Additionally, to quantify the age-dependence patterns across different brain regions in ASD, we examined the age-group patterns of mean brain morphological changes (10 years intervial).

### Case-control Analyses

To estimate the structural brain differences between the ASD and TD groups, we performed the whole-brain analysis for each parameter (CT, SA, and GV) to reveal the ROIs that differed between groups. To reduce inter-individual variation due to head size and inter-image differences etc. caused by voxel-scaling variations, the data of each subject have been normalized to global quantities of the same subject. Specifically, GV are divided by the Estimated Total Intracranial Volume (eTIV) and multiplying 1000; SA are divided by the area of the total white matter in the same hemisphere and multiplying 1000; CT are divided by the mean cortical thickness in the same hemisphere; GC are divided by the mean GC in the same hemisphere; MC are divided by the mean MC in the same hemisphere. After “self-normalization”, z-score is applied to bring all features in the same range. Moreover, the confounding variables are extra variable in principle irrelevant to our analyses, but which can affect the outcome if not properly considered. we then used ComBat (github.com/ Jfortin1/ComBatHarmonization) [57], a batch-effect correction tool, to control for confounding effect of age, sex, and site effects. Finally, the correction for multiple comparisons (Bonferroni), results were considered significant if the Bonferroni-corrected p value was ≤0.05.

### Multi-model Classification with cross-view correlations via low-rank tensor learning

LMFGCN is an intermediate fusion strategy for classification tasks with different types of data (see Fig. 1). The workflow of LMFGCN can be <u>summarized into three components</u>: (1) preprocessing, in which feature normalizations are first performed on each type of data separately in order to minimize inter-individual variation due to differences in head size and inter-image differences etc.; (2) Modality-specific learning through the use of GCNs [32], which involves constructing a weighted sample similarity network for each type of data and subsequently training the GCNs with both the features and the corresponding similarity network for modality-specific learning. And finally, (3) Multi-modal integration via a Low-Rank Multimodal Fusion (LMF) approach [58], which employs a cross-modal discovery tensor calculated from the initial class probability predictions of all modal-specific networks. This tensor is then utilized to train an LMF model, which produces the final predictions. LMF is adept at learning the intra-modality and cross-modal label correlations in the higher-level label space, thereby improving the performance of the model. Of note, LMFGCN is an end-to-end model, with both modality specific GCNs and LMF trained jointly, resulting in a highly integrated and robust model for classification tasks involving different types of data.

**Figure 1.**
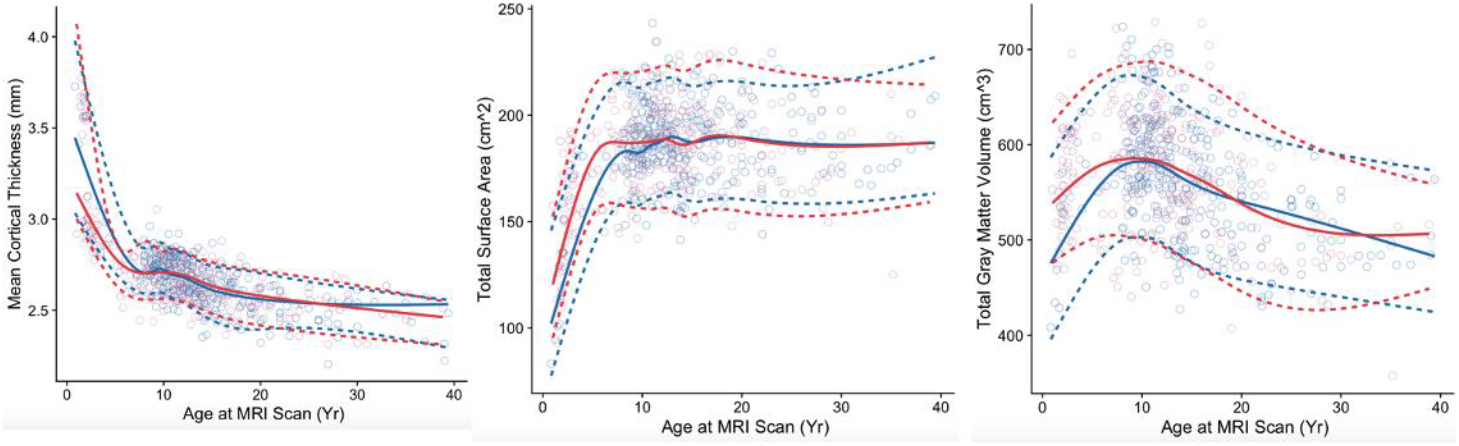
Trajectories of brain morphological alterations across the lifespan. The 5th, 50th, and 95th percentage curves were used to visualize the age-dependent changes for both ASD and TD groups. For mean cortical thickness, TD has a more rapid decline than ASD at the age of 0-8 years. However, this change seems to disappear around the age of 10 years, when growth curves intersect; followed by a near-linear decrease. In contrast, ASD has a larger total brain surface area than TD at the age of 0-8 years, but this rise similarly begins to disappear, after this, no substantial changes in total surface area commonly occur. For total gray matter volume, ASD has a rapid cortical volume larger than TD at age of 0-8 years, peaking at around age 8-10 years, and subsequently decreased from adolescence and adulthood.

### GCNs for feature-specific learning

Previous GCNs models have primarily focused on semi-supervised learning, propagating labels from labeled data to unlabeled ones. However, the applicability of these methods in clinical settings may be limited as the learned GCNs model cannot be directly used to predict new samples when data is not available during the training process [35, 59]. Hence, in this study, we investigate the use of GCNs in supervised learning to capture the intrinsic data structure through graphs during network training. The advantage of using graph NN in a supervised setting lies in its ability to capture both the local intra-class information from multi-modal phenomena within each class and the inter-class information to achieve more discriminative features.

According to conventional definition, a graph can be represented as G = (V, E, W), in which V is the set of nodes, E is the set if edges, and W is the weighted adjacency matrix. In this study, we define each subject as a node. To determine edges, we utilized a cosine similarity measure, which connects each node to its respective neighbors. Further, edges with cosine similarity measure surpassing a predefined threshold value *ϵ*, as determined by parameter *K* = ∑_*i,j*_ *I*(*S*(*x*_*i*_, *x*_*j*_) ≥ *ϵ*)/*n*, have been retained for analysis. Specifically, *A*_*ij*_, which is the adjacency between node *i* and node *j* in the graph, is calculated as:

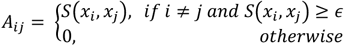

Thus, it is pertinent to note that the GCNs model is predicated on two crucial inputs. First, a feature matrix **X**∈ℝ*n*×*d*, where n represents the number of nodes and d denotes the number of input features. The second input is a description of the graph structure which, in turn, can be aptly illustrated by an adjacency matrix **A**∈ℝ*n*×*n*. The core process of GCNs model is the spectral graph convolution filter which facilitates the convolution operation on non-Euclidean, irregular graph data. Initially, eigen-decomposition is utilized on the normalized Laplacian, which is then followed by the graph Fourier transform (GFT). This conversion step transforms spatial graph convolution into spectral multiplication[60]. Specifically, the normalized Laplacian of graph G is defined as:

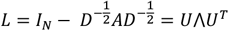

where *I* is the identity matrix, *D* is the degree matrix, A is the adjacency matrix of graph, U is the eigenvectors, and ⋀ is a diagonal matrix of its eigenvalues.

The graph Fourier transform of a signal can be expressed as 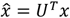. As a results, the convolution on a graph can be defined by multiplying the spectral domain of the signal with a filter as:

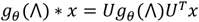

Where *g*_*θ*_ is graph convolution filter [ref], and *x* is the node feature [ref].

To reduce the computation expense and strictly localize the filters, instead of the prior conventional filter, we applied K-order Chebyshev polynomials on normalized Laplacian to compute K localized filters and *g*_*θ*_(⋀) becomes:

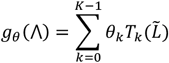

where the parameter *θ* ∈ ℝ^*K*^ is a vector of Chebyshev coefficients,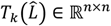 is the Chebyshev polynomial of order k evaluated at 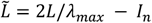.

The cross-entropy loss function was used in our work [ref]:

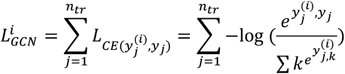

Where *L*_*CE*(·)_ represents the cross-entropy loss function. *y*_*j*_ ∈ ℝ^*c*^ is the one-hot encoded label of the *j*-th training sample and 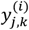 is the *k*-th element in the predicted vector 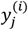 of the *j*-th training sample from the ith data type.

### LMF for multi-modal integration

The utilization of tensor factorization for multimodal data analysis is widely recognized for its exceptional advantages. However, such methods undergo a transformation process of the input tensor, which leads to an exponential augmentation in dimensionality and incurs high computational complexity. This impedes the practicality of the models, particularly when the dataset comprises more than two modalities. Thus, this research endeavors to amalgamate GCNs with a low-rank multimodal fusion approach. This technique enables the learning of both intra-modality and inter-modality information, through the utilization of low-rank weight tensors, leading to effective and efficient multimodal fusion.

In the realm of multimodal fusion, the Low-rank tensor representation technique has proven to be a viable and efficacious method. This technique involves the conversion of multiple input sources into high-dimensional tensors, which are subsequently reduced to a lower dimensional vector space [61]. The tensor representation is derived from the outer product of the input modalities, thus enabling simulation of arbitrary mode subsets. In order to enhance this process, Zadeh et al. [62] introduced an adjustment whereby a value of 1 is added to each representation prior to computing the outer product. So, the input tensor is calculated by representing a single modality:

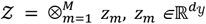

z_m_ is the input representation of an additional 1.

Enter the tensor *𝒵∈*ℝ^*d*^1_,*d*_2_,…,*d*_*m* Through a linear layer *g*(*·*) Produce a vector representation:

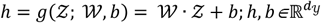

donate *𝒲* as weight, b as bias. So *𝒲* is M +1 order tensor and dimension is *d*_1_ *× d*_2_*×* … *× d*_*m*_*× d*_*h*_.

In the process of performing tensor dot products, we can think of *𝒲* as *d*_*h*_ M-order tensor, that is, it can be divided into 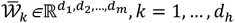, Each 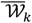 contributes a dimension to the output vector, i.e.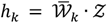.

To address the issue of exponential increase in dimensionality encountered in the tensor-based fusion approach, Liu et al. [58] introduced a LMF method, which *𝒲* decomposes into a set of modality-specific low-rank factors and *𝒵*can also be decomposed into 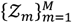. Through this parallel factorization method, LMF can obtain high-dimensional tensors without dominance and directly compute to *h*. Think of *𝒲* as an *d*_*h*_ M-order tensor, each of M-order tensor which can be expressed as 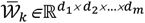, *k* = 1, …, *d*_*h*_, there exists a pattern that is precisely decomposed into vectors:

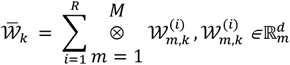

The minimal *R* that makes the decomposition valid is called the rank of the tensor. Based on the 𝒲 and 𝒵 decomposition, we can extrapolate the original calculation formula as follows:

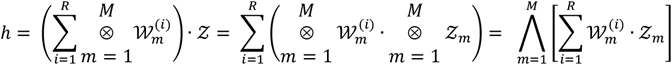

where 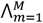 denotes the element-wise product over a sequence of tensors: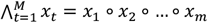. Note that for all m, 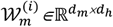 shares the same size for the second dimension.

We can extend Equation 6 for multiple modalities in which we do the element-wise product and summation:

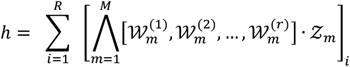

and now the summation is done along the first dimension of the bracketed matrix indicates the *i*-th slice of a matrix. By adopting the proposed approach, the model can be efficiently parameterized with M order-3 tensors, as opposed to implementing sets of vectors as described in Equation 2. This modification not only enhances the overall efficiency but also decouples different modalities in the simplified calculation of h, enabling seamless extension of the approach to any number of modalities.

The structure of the GCNs model pipeline is illustrated in Figure 1. Initially, the GCNs model acquires latent graph representations for each data type. Following this, the LMF model identifies the latent cross-view correlations between the low-rank multi-modalities and enhances the learning performance. The LMF model integrates the initial predictions obtained from different data types, thereby enabling the final prediction derived by LMFGCN to rely on both modality-specific predictions and cross-modality label correlation knowledge learned by LMF. The feature-specific learning aspect of the GCNs is emphasized in LMFGCN. The model is end-to-end, and all networks are trained jointly. During training, each GCNs model is pre-trained for each data type to facilitate good GCNs initialization. Subsequently, during each epoch of the training process, the GCNs model for each data type is first fit to minimize the loss function L. The modal-specific GCNs are then fixed, and LMF is updated to minimize L. The alternation of updating the feature-specific GCNs and LMF continues until the model converges. In addition, 80% of the data is used for training, and 20% is set aside for testing. To avoid overfitting and randomness during training, we used 10-fold cross-validation on training datasets. In each iteration of the scheme, one portion was earmarked as the validation set for model evaluation, and the remaining nine portions were identified as the training set. Furthermore, the 10-fold cross-validation process was stratified to maintain sample percentages for each class in each fold in line with the entire dataset.

### Comparison with different integration methods and other state-of-the-art models

This study investigates a range of classification algorithms, encompassing diverse approaches such as graph filters methods, fusion strategies, and various models. To improve classification performance, we compared GCN-conv and Cheb-conv, which consequently reduced computation expense and localized filters more effectively. Furthermore, we compared the performance of LMFGCN with GCN-conv and Cheb-conv, as well as different fusion strategies, including early, late, and intermediate fusion. Specifically, early fusion directly merges different data types into a large data matrix, which is then inputted into a single model. Late fusion applies unimodal data independently and integrates the results to determine the final consensus. Intermediate fusion learns the latent representations of each modality and models intermodal interactions before executing joint modeling (i.e., LMFGCN). Additionally, we compared the performance of GCNs with state-of-the-art classifiers, such as SVM, XGBoost, LR, RF, DT, and KNN, using accuracy, precision, recall, and F1 value as performance metrics.

## RESULTS

In total, our final sample included 1011 participants with ASD (females = 150, mean 12.7 ± 8.4 years) and 998 TD participants (females = 276, mean 14.6 ± 7.52 years). 470 out of 2,479 participants in the ABIDE I & II and KQJH were rejected. The detailed process of quality assurance please see the supplementary materials. Additionally, distributions of ASD and TD subjects per site were shown in Table. 1. The distribution of age in each site were shown in Figure. S1.

**Table 1.** Demographics of participants included after QA.

| **Demographics** | **TD** | **ASD** | ***p*-Value** |
| --- | --- | --- | --- |
| KQJH | 48 | 216 |  |
| Age, years, mean (SD) | 2.13(0.97) | 2.65(0.83) | <0.001<sup>a</sup> |
| Sex, F/M, n | 23/25 | 38/178 | <0.001<sup>b</sup> |
| ABIDE-I | 481 | 393 |  |
| Age, years, mean (SD) | 13.65(6.69) | 13.72(6.38) | 0.873<sup>a</sup> |
| Sex, F/M, n | 163/318 | 58/335 | .0.001<sup>b</sup> |
| ABIDE-II | 469 | 402 |  |
| Age, years, mean (SD) | 16.85(7.23) | 17.04(7.95) | 0.705<sup>a</sup> |
| Sex, F/M, n | 90/379 | 54/348 | 0.023<sup>b</sup> |
| **Total** | 998 | 1011 |  |
| Age, years, mean (SD) | 14.6(7.52) | 12.7(8.4) | <0.001<sup>a</sup> |
| Sex, F/M, n | 276/722 | 150/861 | <0.001<sup>b</sup> |

ABIDE, Autism Brain Imaging Data Exchange; TD, typical development; ASD, autism spectrum disorder; F, female; M, male.
<sup>a</sup>*p*-Value was calculated using a two-sample *t*-test.
<sup>b</sup>*p*-Value was calculated using $\chi^2$ test.

### Tractography analysis

The growth trajectories of MCT, TGV, and TSA were explicitly estimated using GAMLSS (Figure 1), providing formal quantification of developmental changes in between-subject variability. To visualize the age-dependent changes for both ASD and TD groups, the 5th, 50th, and 95th percentage curves were utilized. Corresponding AIC and BIC for each model in the training set are presented in Table S1-6. In both groups, age-related cortical thinning was observed across the lifespan, although ASD demonstrated thinner cortical thickness than TD during the age range of 0-8 years. This discrepancy appears to diminish around the age of 8-10 years, when growth curves intersect and is subsequently followed by a near-linear decrease. Conversely, ASD demonstrated a larger brain area than TD during the same age range. However, after growth curves intersect, both curves experienced an arrest in growth. For TGV, ASD displayed a rapid cortical volume increase larger than TD during the age range of 0-8 years, peaking at approximately 8-10 years of age, and subsequently decreasing from adolescence to adulthood. Age effects appeared to be slightly non-linear, while the distribution was mildly heteroskedastic and non-gaussian.

Furthermore, the findings provide compelling evidence for distinct developmental trajectories in various brain regions of individuals with ASD. Specifically, as illustrated in Figure. S1-3, the cortical thickness-surface area exhibited alterations throughout different age groups, indicating a separable course of development that contributes to changes in gray matter volume. In addition, the study supports the notion that alterations of CT and GV in ASD demonstrate diverse temporal and regional patterns across different age groups and brain regions. Notably, these patterns appear to vary depending on the regional sequences of typical early brain development (i.e., from back to front).

### Case-Control Differences

A series of general linear models were utilized to compare CT, SA, and GV between individuals with ASD and TD individuals. The statistical analysis revealed significant differences in cortical measures between the two groups, as illustrated in Figure 2/Table S7. Specifically, individuals with ASD exhibited thinner CT in the left precentral, right postcentral, paracentral, caudal middle frontal and opercularis, as well as bilateral parahippocampal regions. Conversely, thicker CT was observed in the left medial orbitofrontal, rostral middle frontal, right lingual, and bilateral lateral occipital regions (Figure 2A). In regard to SA, individuals with ASD demonstrated decreased SA in the right medial orbitofrontal, and bilateral rostral anterior cingulate and lateral orbitofrontal regions, while concurrently exhibiting increased SA in the left isthmus cingulate, transverse temporal, and lateral occipital regions, as well as bilateral precuneus regions (Figure 2B). Finally, with regards to GV, larger GV was found in the left medial orbitofrontal and isthmus cingulate, right precuneus, and bilateral lateral occipital regions in individuals with ASD, while smaller GV was observed in the right precentral and caudal anterior cingulate regions, as well as the bilateral rostral anterior cingulate and caudal middle frontal regions (Figure 2C).

**Figure 2.**
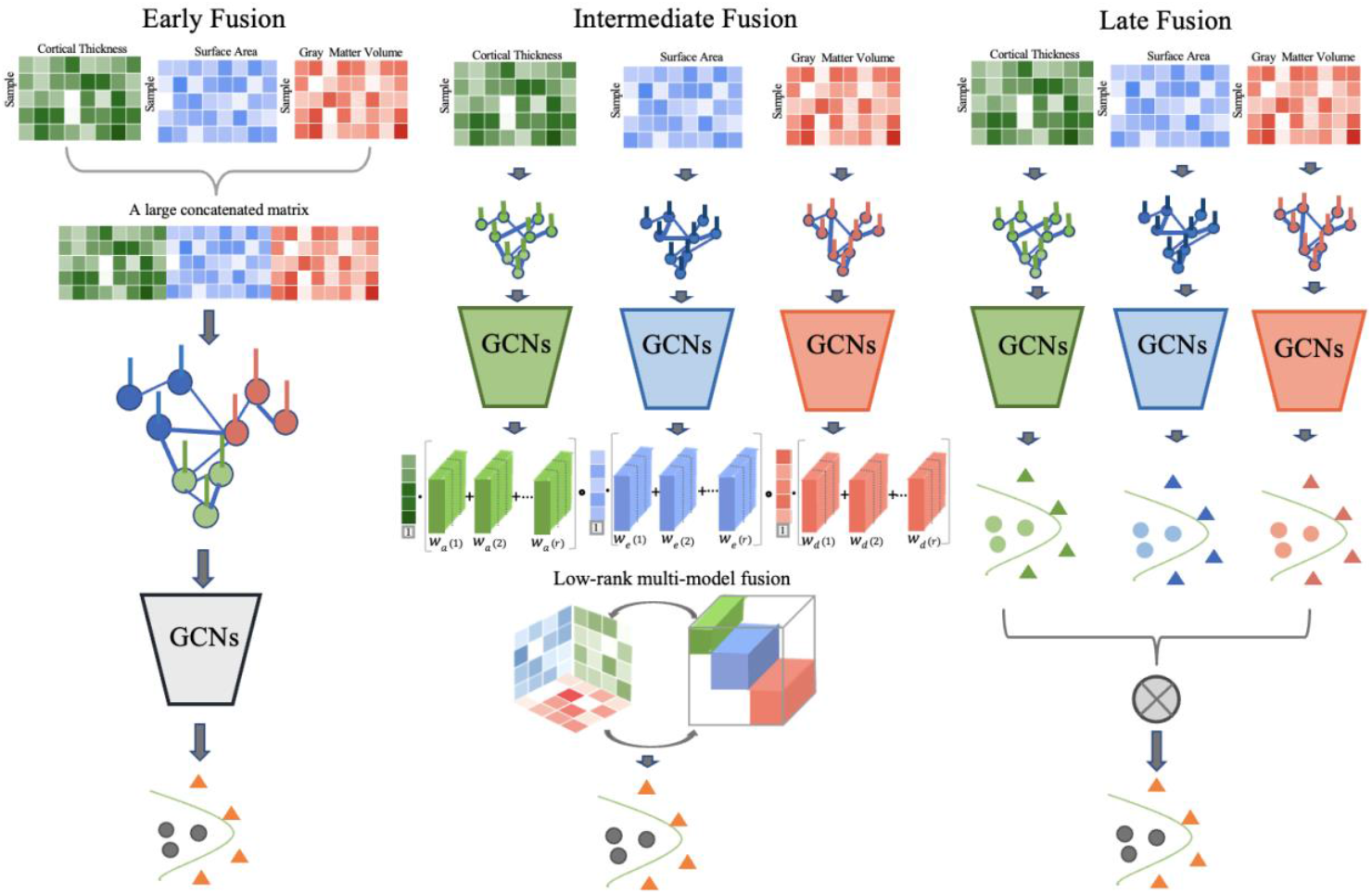
Overview of different fusion strategies for multimodal data. Early fusion model (Left panel) concatenates original features at the input level. Intermediate fusion (Middle panel) not only extracts the intra-modality information in each mode, but also captures the correlation between modes. Late fusion (right panel) aggregates predictions at the decision level.

**Figure 3.**
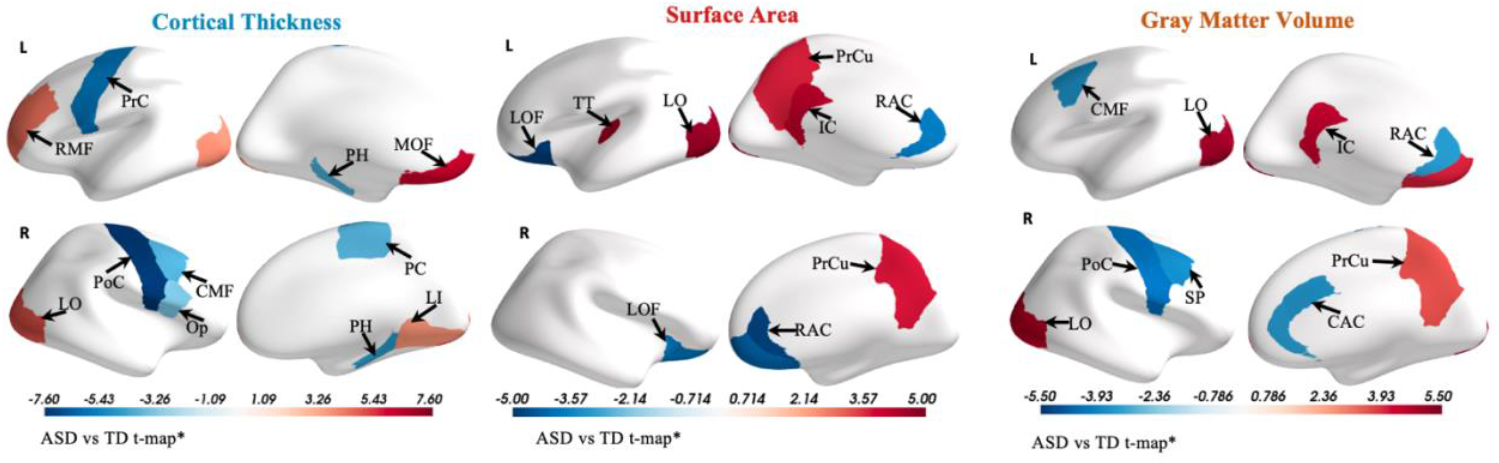
Cortical alteration differences between ASD and TD. Surface maps show of the cortical regions for which there were significant group differences in cortical thickness, surface area, and gray matter volume after the Bonferroni-correction. Abbreviations: Precentral (PrC), Paracentral (PC), Postcentral (PoC), Precuneus (PrCu), Lateral occipital (LO), lateral orbitofrontal (LOF), Rostral middle frontal (RMF), Pars opercularis (Op), Parahippocampal (PH), Transverse temporal (TT), Lingual (LI), caudal anterior cingulate (CAC), Rostral anterior cingulate (RAC), caudal middle frontal (CMF), Isthmus occipital (IC).

### Classification performance

In this study, we utilized a combination of GCM and LMF methods to perform classification of ASD based on various feature types. Firstly, a weighted sample similarity network was constructed for each feature type using cosine similarity. Subsequently, these features and the respective similarity networks were incorporated into the corresponding GCN models to generate initial predictions. These initial predictions were then reshaped into a vector and processed through low-rank multi-model fusion to yield final label predictions. Performance evaluation of our LMFGCN approach involved comparing graph filter methods, fusion strategies, and several machine/deep learning methods for ASD classification. We employed ten-fold cross-validation to fit all methods and evaluated the resulting models on independent test data, with the mean and standard deviation of evaluation metrics being reported (refer to Figure S3). Our results demonstrate that the LMFGCN approach based on Cheb-conv outperforms other approaches, yielding a highest accuracy of approximately 81.77% on the test dataset (please see Table 2 for further details).

**Table 2.** Comparison with early and late fusion model and traditional machine learning models. LMFGCN classifier achieved the best performance due to its ability to integrate inter-modality and intra-modality information according to their latent representation for the final predictions.

| Model | Accuracy | Precision | Recall | F1 |
| --- | --- | --- | --- | --- |
| XGBoost | 0.5821 | 0.5829 | 0.5823 | 0.5814 |
| KNN | 0.5945 | 0.5999 | 0.5951 | 0.5898 |
| DT | 0.5697 | 0.5702 | 0.5698 | 0.5692 |
| RF | 0.5945 | 0.5999 | 0.5951 | 0.5898 |
| SVM | 0.5995 | 0.6124 | 0.6003 | 0.5888 |
| LR | 0.5995 | 0.6025 | 0.5999 | 0.5971 |
| Early-Fusion (GCNConv) | 0.7403 | 0.7414 | 0.74 | 0.7398 |
| Late-Fusion (GCNConv) | 0.7348 | 0.7396 | 0.7392 | 0.7346 |
| Intermediate-Fusion (GCNConv) | 0.7289 | 0.7443 | 0.7313 | 0.7257 |
| **LMFGCN(ChebConv)** | **0.8177** | **0.8215** | **0.8187** | **0.8174** |

Abbreviations: Support vector machine classifier (SVM), Linear regression (LR), Random Forest (RF), and K-nearest neighbor (KNN), Decision Tree (DT).

## Discussion

Brain development in individuals with ASD is a dynamic and nonlinear process, particularly during early stage [63-65]. Abnormal development at the critical periods may lead to significant impairments in the establishment of neural circuitry that underlie cognitive, socio-emotional, and communication functions[5, 9]. In the current study, we adopted the GAMLSS model to qualify normative trajectories for cortical morphological measures in the largest sample to date of patients with ASD and TD across the lifespan. Our findings indicate that cortical thickness-surface area trajectories are separate and distinct across the lifespan. We also identified crucial neural biomarkers linked to ASD, which may have implications for early detection and intervention. Additionally, we utilized a deep-learning framework called LMFGCN, which integrates multiple types of cerebral cortical features to classify ASD. Our approach outperformed other methods tested on a large dataset across multiple sites, achieving classification accuracy exceeding 81.77%.

The proper maturation of the brain during the early stages of neurodevelopment is critical for understanding the pathology of ASD [66, 67]. In this critical period, structural and functional abnormalities in the brain can impair the development of social, emotional, and cognitive functions, resulting in significant and far-reaching consequences later in life [68, 69]. The results of our investigation indicate that in the initial years of development, MCT, TSA, and TGV display swift and dynamic trajectories that gradually diminish around 8-10 years of age. These results are consistent with the largest existing imaging study of children with ASD [13, 25]. It is noteworthy, however, that after the intersection of the growth curve (around age 10 years), there appears to be a period of “pseudonormalization” that may not reflect broader structural or functional normalizations. These pseudonormalizations in neurodevelopmental trajectories may be related to underlying neural complexity and contribute to the heterogeneity observed in ASD. Moreover, our study demonstrated that the larger brain volume observed in very young children with ASD is predominantly attributed to increased SA, rather than CT. This aligns with similar findings on increased total surface area at younger ages [19, 70] and decreasing CT with age in ASD [25]. Our research also supports the notion of divergent developmental trajectories of cortical thickness and surface area throughout the lifespan [19, 71], which suggests that distinct mature trajectories for both measures are related to different aspects of cortical neuropathology and represent unique features of the cortical architecture. The early expansion of cortical surface area was previously thought to result from symmetrical proliferation of neuronal progenitor cells, while later events in development determine the number of neurons and synapses per column, which are reflected in cortical thickness according to the radial unit hypothesis of Rakic [24, 72, 73]. Therefore, the finding of early enlargement of the brain in ASD, caused by an accelerated expansion of SA but not CT, is of significant interest as it points towards specific genetic and neurobiological mechanisms that may be impaired in ASD. This highlights the need for the development of innovative neuroimaging measurements that offer a higher degree of specificity with regard to particular underlying mechanisms. The development of brain structure is a dynamic process, and it has been suggested that atypical regional alterations in ASD may contribute to the manifestation of relevant autism traits. Although the vast previous findings[26, 69, 74-76] have revealed cortical abnormalities in individual with ASD of varying ages and has indicated that these changes may be associated with autism-related symptoms, the results have not been entirely inconsistent. Accordingly, we systematically investigated brain morphometry differences between ASD and TD in a large-scale study across the lifespan. Our results indicated that individuals with ASD demonstrated the greatest decreases in both CT and GV in the sensory-motor cortex and parahippocampal cortex, whereas they exhibited significant increases in the left orbitofrontal cortex. Such findings may highlight disruptions in cognition, movement, and memory formation as neural correlates of ASD, ultimately leading to impaired social perception and communication skills [25, 77]. The left orbitofrontal cortex, a key region for social cognition and repetitive behaviors, was consistently shown to be associated with ASD [78, 79]. Conversely, individuals with ASD demonstrated decreased SA and GV in the anterior cingulate cortex, which may contribute to abnormal reciprocal social interaction [80]. Additionally, increased left transverse temporal, precuneus cortex, and both posterior cingulate were found to be associated with auditory processing and self-referential processing in ASD [81-84]. Interestingly, a common increase in the left lateral occipital cortex was observed for all cortical measures, which may play a role in face perception and has been linked to communication deficits in boys with ASD [85, 86]. Our findings reveal distinct patterns of cortical changes in CT and SA between individuals with ASD and TD, and their contribution to GV changes across the lifespan suggesting separate developmental trajectories and unique genetic determinants [69]. Moreover, the abnormal cortical patterns for CT and SA were largely nonoverlapping, which further supported their separatable developmental trajectories and indicate that they have distinct genetic determinants.

Although a previous study has shown the effectiveness of GCNs in distinguishing individuals with ASD from TD, our study presents several unique advantages. Firstly, while the previous GCN models primarily employed a semi-supervised learning approach, we propose a supervised GCN model based on a graph similarity integration strategy. This implies that our model is highly flexible and can generalize well to new data. Secondly, previous data fusion methods often suffer from exponentially increasing dimensions and computational complexity. In contrast, our model combines GCNs for multi-modality-specific learning and LMF for multi-model integration. This enables us to transform a high-dimensional tensor into a lower-dimensional vector space that has the potential to improve the model’s performance. Thirdly, we used ten-fold cross-validation to evaluate our GCNs model and compared it to a set of other methods in a larger sample size. This helps to reduce the risk of overfitting the training dataset and provides more reliable and generalizable results. Indeed, our results demonstrated that LMFGCN exhibited superior prediction performance compared to other common classifiers.

There are several limitations to the current study. Firstly, current study has a larger sample size cross the life span, but various participating sites and sex differences still be challenging for multi-sits datasets. While we made effects to control for these effects in our models, it cannot fully exclude potential influence. Secondly, the classification model for ASD must be replicated on other independent datasets before any application in clinical decision-making. Lastly, the developmental trajectories for brain changes in ASD are predominantly drawn from our current cross-sectional data, future longitudinal studies with larger sample sizes are essential to clarify reported developmental trajectories and determine whether distinct early developmental trajectories can predict brain changes in the later life.

## Conclusion

In conclusion, our study offers a comprehensive perspective on the cortical development of the brain and elucidates a rapid and dynamic structural transformation during early stages of life, which is a pivotal period for comprehending the neurobiology of ASD. The altered patterns for CT and SA are congruent with those in GV, which signify diverse facets of the underlying neural architecture. Additionally, LMFGCN, which incorporates both GCNs and LMF models, achieved noteworthy outcomes on a large multi-site dataset, suggesting the potential of LMFGCN in characterizing ASD. These findings hold significant implications for understanding the neuropathology of ASD and offer a promising avenue for utilizing GCNs in ASD classification.

## Acknowledgments

This study was supported by the National Natural Science Foundation of China (No. U22A2041), Shenzhen Science and Technology Program (No. KQTD20200820113106007), China Postdoctoral Science Foundation (No. 2021M703375), National Science Foundation of China (No. 62272449), Strategic Priority CAS Project (No. XDB38050100), the Key Research and Development Project of Guangdong Province (Grant No. 2021B0101310002), the Shenzhen Basic Research Fund (No. RCYX2020071411473419), and Youth Innovation Promotion Association (No. Y2021101). Data used in this study were obtained from the Autism Brain Imaging Data Exchange (http://fcon_1000.projects.nitrc.org/indi/abide/) and We would also like to thank the year-by-year brain templates (1-6 Years) shared by Zhi Yang (https://osf.io/fm7cq/?view_only=9716e89f09e04b4bb2b4f0323ab2b684).

## Availability of data and code

The datasets supporting the conclusions of this article are available in the Autism Brain Imaging Data Exchange and National Database for Autism Research repositories; (ABIDE), http://fcon_1000.projects.nitrc.org/indi/abide/.

All code of GAMSS and LMFGCN can be found at https://github.com/ASD-trajectory-and-classification.

## Author contributions

YL.W. contributed to the conception and design of the study. YL.W. performed and wrote the paper. S.T., SQ.M., and RM.M. participated in the discussion of methods and results. YJ.W. and Y.P. reviewed the manuscript and provided comments. All authors have read and approved the final version of the submitted manuscript.

