## Supplementary material for "Distinct Neurodevelopmental Patterns and Intermediate Integration-Based Predictive Modeling in Autism Spectral Disorder": Supplemantary: Supplementary.pdf

### Supplementary Materials:

### MRI acquisition

All T1-weighted scans were chosen from 1112 ABIDE-I participants (539 ASD participants and 573 TC), 1114 ABIDE-II participants (521 ASD participants and 593 TC participants), and 323 KQJH participants (237 ASD participants, 86 TC participants). A detailed introduction of the participants and scan parameters in each center for ABIDE dataset (ABIDE-I [1] and ABIDE-II [2]) is available at http://fcon_1000.projects.nitrc.org/indi/abide/. KQJH dataset was acquired in the sagittal plane on 3 T scanners (Siemens Skyra) using a magnetization prepared rapid gradient echo (MPRAGE) sequence. Acquisition parameters were as follows: TR/TE/TI = 2020/2.11/900 ms; flip angle: 8°; FOV = 224 × 224 $mm^2$; and 1 × 1 × 1 mm3 isotropic voxel. Before scanning, the sedation was performed with 10% chloral hydrate (50 mg/ml) orally or by enema to reduce the influence of children's head movement on image quality.

### Quality assurance (QA)

The quality control was assessed by inspecting the image and the cortical reconstructed results across individual brains by two independent reviewers (YL.W. and S.T.). To enhance the accuracy of QA and to investigate the reason for exclusion, we use quality control software for MRI data (MRIQC [3] ) to compute image quality metrics for each dataset (see Table. S8/9/10).

In ABIDE-I dataset, out of the 238 inaccurately segmented scans, 116 had abnormally low contrast-to-noise ratio (CNR), and the remaining 122 scans showed evidence of high subject motion, as estimated using the entropy focus criterion (EFC). For these reasons, the data from these 238 participants were excluded from further analyses, resulting in a final sample size of N = 874 subjects.

In ABIDE-II dataset, out of the 243 inaccurately segmented scans, 63 had abnormally low CNR, and the remaining 180 scans showed evidence of high subject motion, as estimated using the EFC. For these reasons, the data from these 243 participants were excluded from further analyses, resulting in a final sample size of N = 871 subjects.

KQJH dataset were subjected to the same quality assurance steps as ABIDE-I & II. out of the 59 inaccurately segmented scans, 32 had abnormally low CNR, and the remaining 27 scans showed evidence of high subject motion, as estimated using the EFC. For these reasons, the data from these 107 participants were excluded from further analyses, resulting in a final sample size of N = 216 subjects.

After the QA steps described above, ABIDE I consisted of 393 ASD participants (N=58 female), age range 7-64 (M = 12.72, SD = 6.38) and 481 TD participants (163 female) age range 6-56 (M = 12.65, SD = 6.69). ABIDE II consisted of 402 ASD participants (N=54 female), age range 7-64 (M = 17.04, SD = 7.95) and 469 TD participants (90 female) age range 6-56 (M = 16.85, SD = 7.23). KQJH consisted of 216 ASD participants (N=38 female), age range 1-6 (M = 2.65, SD = 0.83) and 48 TD participants (23 female) age range 0-6 (M = 2.13, SD = 0.97). Moreover, age at time of scan in years per collection (sites) was plotted in Figure S1.

### Image preprocessing and feature extraction

**For 0-2 Years:**

The T1-weighted images of participants between 0-2 years of age were analyzed using Infant Freesurfer software (https://surfer.nmr.mgh.harvard.edu/fswiki/infantFS). Infant Freesurfer is a pipeline of automated segmentation and surface extraction based on Freesurfer for T1-weighted neuroimaging data for infants [4]. Automatic processing steps include intensity normalization, skull stripping, and segmentation of the cortex, white matter, and subcortical structures. Segmentation involves using a multi-atlas approach in which multiple atlases are first registered to subject space and the labels are transferred. The atlases were developed from infant MRI scans [5]. To create the atlas, manually segmented labels were developed using MRI scans from a representative sample of infants (0–2 years of age). In the current study, developmentally appropriate atlases for 4- and 6-month-old infants were employed. A total of 68 regionally distributed measurements (grey matter volume, surface area, and cortical thickness) were extracted from the “aparc” [6] in each subject.

**For 3-6 Years:**

The T1-weighted images of participants between 3-6 years of age were estimated using FreeSurfer (version 7.1.117) and ANTs (version 2.0). Since the brain size and gray-white matter contrast change with age during childhood periods, we first applied the age-specific brain templates [7] and created prior probabilistic tissue maps/masks including gray matter, white matter, cerebrospinal fluid (CSF), subcortical structures, brain stem, and cerebellum (see Data and Code availability). Then, all images were transformed and inputted to the FreeSurfer "recon-all" pipeline, and the corresponding white matter and brain extraction masks obtained above were injected into the pipeline for tissue segmentation. The detailed information of MRI preprocessing process sees the blow descriptions (over 6 Years).

**Over 6 Years:**

Cortical reconstruction and volumetric segmentation for all participants were performed with the FreeSurfer software, version 7.1.117 (http://surfer.nmr.mgh.harvard.edu/). It begins with transformation to Talaraich space, intensity inhomogeneity correction, bias field correction [8], and skull-stripping [9]. Thereafter, WM is separated from GM and other tissues and the volume within the created WM–GM boundary is filled. After this, the surface is tessellated and smoothed. After these preprocessing steps are completed, the surface is inflated [10] and registered to a spherical atlas. This method adapts to the folding pattern of each individual brain, utilizing consistent folding patterns such as the central sulcus and the sylvian fissure as landmarks, allowing for high localization accuracy. FreeSurfer uses probabilistic approach based on Markov random fields for automated labeling of brain regions. Cortical thickness is calculated as the average distance between the WM–GM boundary and the pial surface on the tessellated surface. The FreeSurfer algorithm automatically parcellates the cortex and assigns a neuroanatomical label to each location on a cortical surface model based on probabilistic information. The parcellation scheme of the Desikan–Killiany atlas [6, 11] was used to divide the cortex into 34 regions per hemisphere.

**Table. S1**. MCT quality of a model fit in the train set in TD.

| **Model** | **AIC** | **ΔAIC** | **AIC weights** | **BIC** | **ΔBIC** | **BIC weights** |
|---|---|---|---|---|---|---|
| m1a | -1176.384 | 0 | 5.399 | -1096.167 | 9.999 | 9.999 |
| m1b | -1176.460 | 0 | 5.608 | -1078.056 | 1.167 | 1.167 |
| m1c | -1194.451 | 0.452 | 4.523 | -1070.866 | 3.205 | 3.205 |
| m1d | -1189.003 | 0.03 | 2.968 | -1045.918 | 1.226 | 1.226 |
| m1e | -1194.722 | 0.518 | 5.179 | -1067.457 | **5.828** | **5.828** |

**Table. S2**. MCT quality of a model fit in the train set in ASD.

| **Model** | **AIC** | **ΔAIC** | **AIC weights** | **BIC** | **ΔBIC** | **BIC weights** |
|---|---|---|---|---|---|---|
| m1a | -1013.294 | 0 | 2.216 | -921.794 | 0.017 | 0.017 |
| m1b | -1019.747 | 0 | 5.583 | -910.376 | 0 | 0 |
| m1c | -1042.971 | 0.617 | 6.165 | -928.077 | 0.248 | 0.248 |
| m1d | -1037.891 | 0.049 | 4.863 | -899.374 | 0 | 0 |
| m1e | -1041.751 | 0.335 | 3.349 | -930.264 | **0.741** | **0.741** |

**Table. S3**. TSA quality of a model fit in the train set in TD.

| **Model** | **AIC** | **ΔAIC** | **AIC weights** | **BIC** | **ΔBIC** | **BIC weights** |
|---|---|---|---|---|---|---|
| m1a | 5917.143 | 0 | 0 | 5981.542 | 0 | 0 |
| m1b | 5797.215 | 0.179 | 0.179 | 5881.842 | 0.003 | 0.003 |
| m1c | 5794.407 | 0729 | 0729 | 5893.745 | 0 | 0 |
| m1d | 5798.538 | 0.092 | 0.092 | 5915.035 | 0 | 0 |
| m1e | 5812.53 | 0 | 0 | 5917.205 | **0** | **0** |

**Table. S4.** TSA quality of a model fit in the train set in ASD.

| Model | AIC | ΔAIC | AIC weights | BIC | ΔBIC | BIC weights |
|---|---|---|---|---|---|---|
| m1a | 6178.629 | 0 | 0 | 6243.123 | 0 | 0 |
| m1b | 6088.446 | 0.018 | 0.018 | 6165.992 | 0.999 | 0.999 |
| m1c | 6091.601 | 0.002 | 0.002 | 6180.681 | 0 | 0 |
| m1d | 6079.592 | 0.986 | 0.986 | 6193.296 | 0 | 0 |
| m1e | 6114.621 | 0 | 0 | 6228.230 | **0** | **0** |

**Table. S5**. TGV quality of a model fit in the train set in TD.

| Model | AIC | ΔAIC | AIC weights | BIC | ΔBIC | BIC weights |
|---|---|---|---|---|---|---|
| m1a | 7513.614 | 0 | 0 | 7585.569 | 0 | 0 |
| m1b | 7413.831 | 0 | 0 | 7486.466 | 0.002 | 0.002 |
| m1c | 7506.856 | 0 | 0 | 7580.772 | 0 | 0 |
| m1d | 7399.117 | 0.507 | 0.507 | 7485.263 | 0.003 | 0.003 |
| m1e | 7399.172 | 0.493 | **0.493** | 7473.871 | **0.995** | **0.995** |

**Table. S6**. TGV quality of a model fit in the train set in ASD.

| Model | AIC | ΔAIC | AIC weights | BIC | ΔBIC | BIC weights |
|---|---|---|---|---|---|---|
| m1a | 7664.468 | 0 | 0 | 7730.337 | 0 | 0 |
| m1b | 7599.203 | 0.103 | 0.103 | 7671.842 | 0.198 | 0.198 |
| m1c | 7663.216 | 0 | 0 | 7737.914 | 0 | 0 |
| m1d | 7597.303 | 0.265 | 0.265 | 7678.852 | 0.006 | 0.006 |
| m1e | 7595.564 | 0.632 | **0.632** | 7669.058 | **0.796** | **0.796** |

**Table. S7**. The results of the whole-brain analysis in brain morphological parameters (ASD vs. TD groups). All p-values are Holm-Bonferroni corrected (< 0.05).

| Brain region | Hemisphere | Contrast | T-value | *p*-value |
|---|---|---|---|---|
| **Gray Matter Volume** | | | | |
| Medial orbitofrontal | L | ASD>TD | 3.96 | 0.005 |
| Isthmus cingulate | L | ASD>TD | 4.21 | 0.002 |
| Lateral occipital | L | ASD>TD | 5.12 | <0.001 |
| Rostral anterior cingulate | L | ASD<TD | -3.49 | 0.034 |
| Caudal middle frontal | L | ASD<TD | -3.57 | 0.0245 |
| Precuneus | R | ASD>TD | 3.51 | 0.031 |
| Lateral occipital | R | ASD>TD | 4.51 | <0.001 |
| Precentral | R | ASD<TD | -4.38 | <0.001 |
| Caudal anterior cingulate | R | ASD<TD | -3.59 | 0.023 |
| Rostral anterior cingulate | R | ASD<TD | -3.44 | 0.04 |
| Caudal middle frontal | R | ASD<TD | -3.87 | 0.007 |
| **Surface Area** | | | | |
| Isthmus cingulate | L | ASD>TD | 4.13 | 0.003 |
| Transverse temporal | L | ASD>TD | 4.51 | <0.001 |
| Lateral occipital | L | ASD>TD | 7.74 | <0.001 |
| Precuneus | L | ASD>TD | 3.7 | 0.015 |
| Rostral anterior cingulate | L | ASD<TD | -3.84 | 0.009 |
| Lateral orbitofrontal | L | ASD<TD | -4.86 | <0.001 |
| precuneus | R | ASD>TD | 3.7 | 0.015 |
| Rostral anterior cingulate | R | ASD<TD | -4.7 | <0.001 |
| Lateral orbitofrontal | R | ASD<TD | -3.81 | 0.01 |

| | | | | |
|---|---|---|---|---|
| Medial orbitofrontal | R | ASD<TD | -4.21 | 0.002 |
| **Cortical Thickness** | | | | |
| Medial orbitofrontal | L | ASD>TD | 6.14 | <0.001 |
| Rostral middle frontal | L | ASD>TD | 4.17 | 0.002 |
| Lateral occipital regions | L | ASD>TD | 3.71 | 0.015 |
| Precentral | L | ASD<TD | -5.82 | <0.001 |
| Parahippocampal | L | ASD<TD | -4.16 | 0.002 |
| Caudal middle frontal | R | ASD<TD | -4.13 | 0.003 |
| Opercularis | R | ASD<TD | -3.39 | <0.049 |
| Precentral | R | ASD<TD | -7.58 | <0.001 |
| Paracentral | R | ASD<TD | -4.08 | 0.003 |
| Parahippocampal | R | ASD<TD | -4.65 | <0.001 |
| Lingual | R | ASD>TD | -3.50 | 0.033 |
| Lateral occipital | R | ASD>TD | 4.71 | <0.001 |

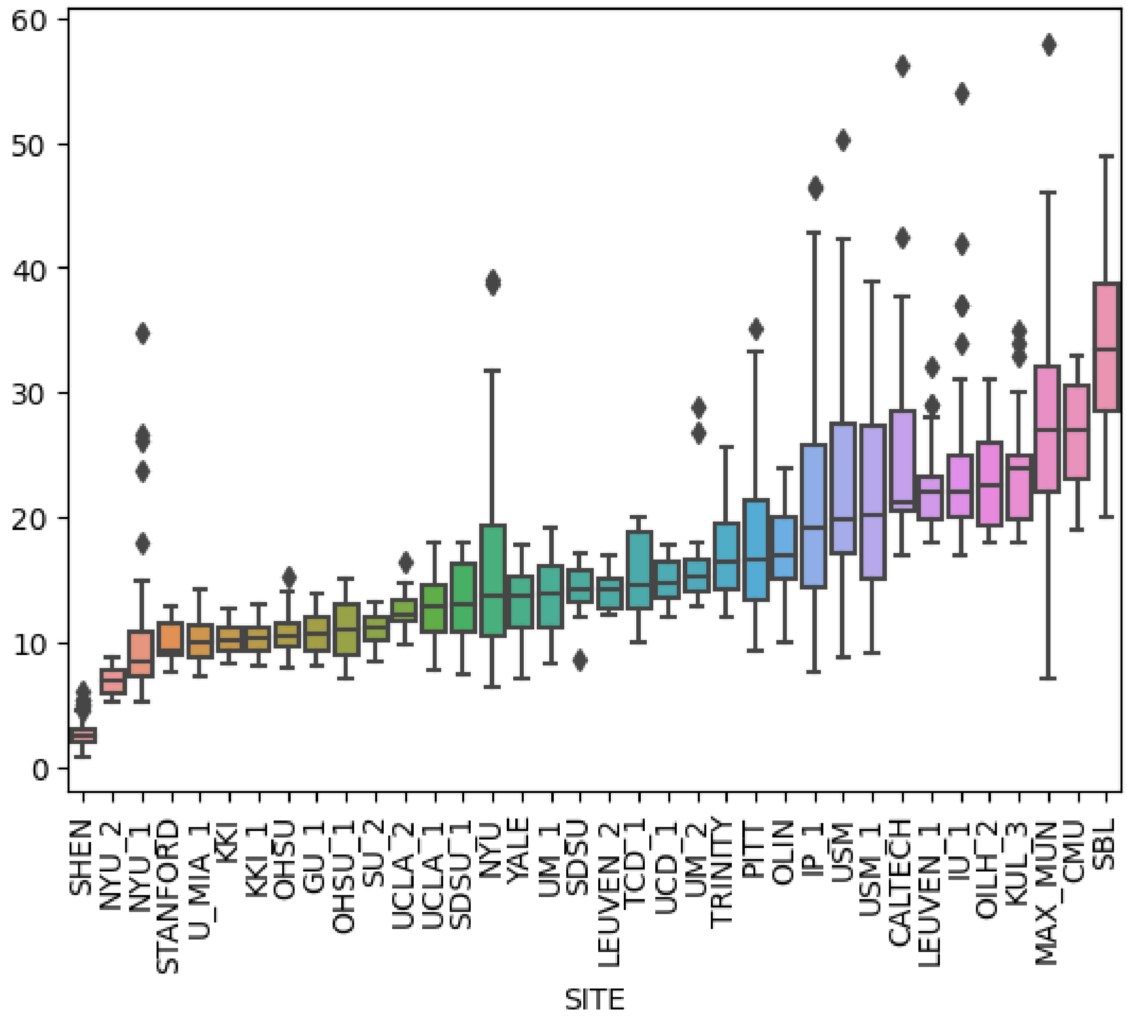


**Figure S1.** Age at time of scan in years per collection (ordered by mean age per collection), irrespective of diagnostic group.

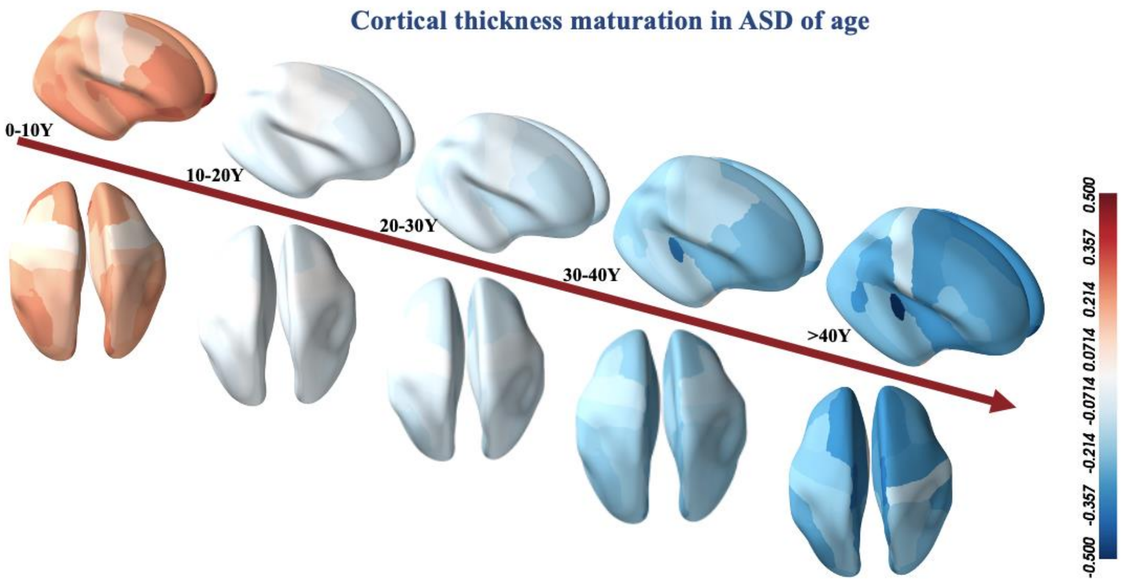


**Figure S2.** Dynamic maps of mean cortical thickness in ASD groups across the lifespan (the interval is ten years). The colors indicate the changes of cortical thickness across ages. Most cortical regions become thinner from birth to older years old.

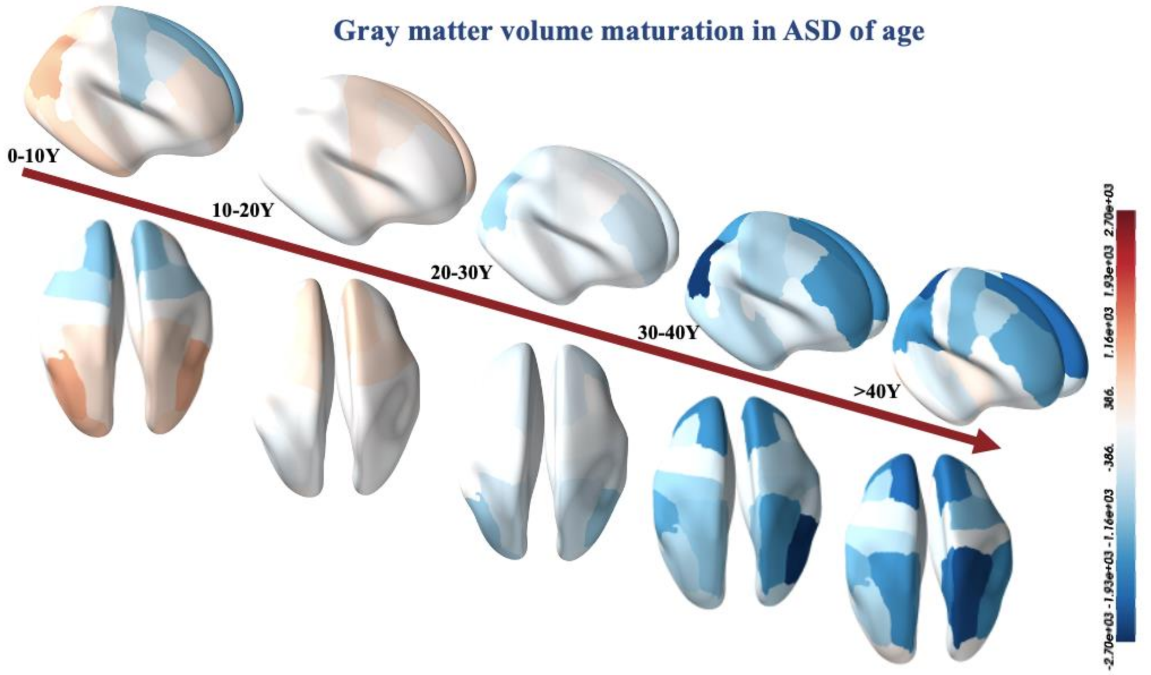


**Figure S3.** Dynamic maps of total surface area in ASD groups across the lifespan (the interval is ten years). The colors indicate the changes of cortical thickness across ages. Most cortical regions become thinner from birth to older years old.

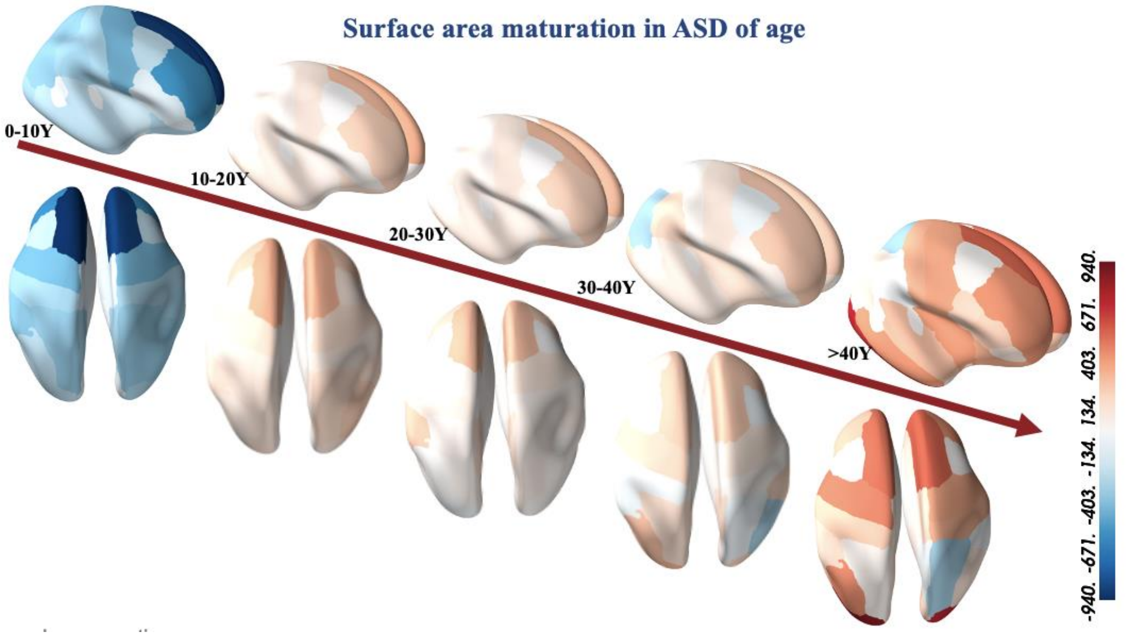


**Figure S4.** Dynamic maps of total gray matter volume in ASD groups across the lifespan (the interval is ten years). The colors indicate changes of cortical thickness across ages. Most cortical regions become thinner from birth to older years old.

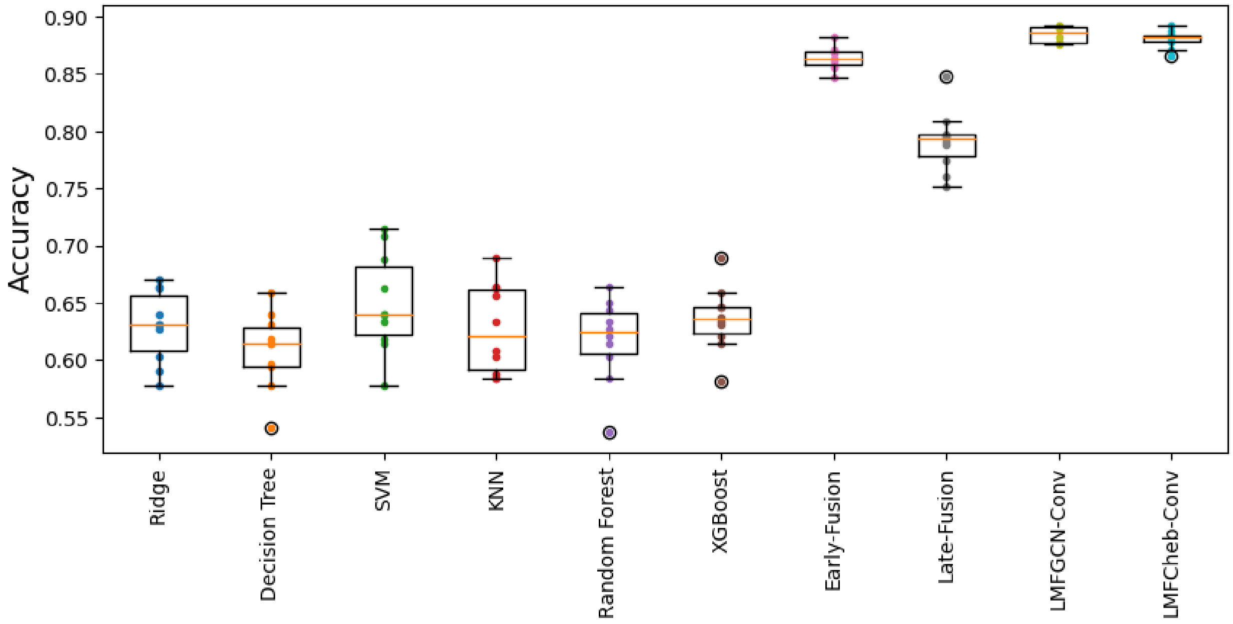


**Figure S5** 10-fold cross-validation of LMFGCN performance in comparison to various filter strategies, integration methods, and classical machine learning models. Boxplots summarize accuracy scores obtained across CV-folds.
